# EF-G Mutations Reveal Correlation between Power Stroke and Translocation Fidelity in Protein Synthesis

**DOI:** 10.1101/2025.06.24.661293

**Authors:** Yanjun Chen, Jacob H. Steele, Shoujun Xu, Yuhong Wang

## Abstract

One of the most essential biomolecular functions is the ribosome translocation for protein synthesis, in which the ribosome moves on the mRNA by primarily three nucleotides per step in the presence of elongation factor G (EF-G). The large conformational changes of EF-G generate significant mechanical force, referred to as power stroke. Quantification of power stroke remains under debate and its correlation with translocation fidelity has not been observed. In this work, we use quantum sensing techniques for measuring both the EF-G power stroke and its influence on ribosome translocation steps. Two EF-G mutants, H584K and Q508K, were expressed, with the mutated residues directly interacting with tRNA. H584K, which interacts on codon-anticodon minihelix, produced less power stroke of 60±6 pN and induced “-1” frameshifting, wherein the ribosome translocated only two nucleotides. In contrast, Q508K, which interacts with tRNA residue 37 immediately outside the codon-anticodon minihelix, exhibited a normal power stroke of 89±11 pN and maintained canonical 3-nucleotide translocation. These findings provide direct mechanistic evidence that the pivotal point and EF-G power stroke are critical for maintaining translocation fidelity and highlight the potential of quantum sensing for chemical biology.

## INTRODUCTION

The ribosome is a molecular machine that translates genetic codes in mRNA into proteins. Its function involves many mechanical parameters, such as displacement, rotation, ratcheting, and power stroke.^1–3^ It is critical for the ribosomes to translocate on the messenger RNA (mRNA) by exactly three nucleotides (nts) per step to produce the correct protein sequence.^4, 5^ In this complex process, mRNA threads through the two subunits of the ribosome, the 50S and 30S subunits in bacteria. The E, P, and A sites are initially aligned, and the transfer RNA (tRNA) is bound at the A site. EF-G, in complex with guanosine triphosphate (GTP), accelerates and stabilizes the ribosome’s twisting motions while itself undergoes a large conformational change driven by GTP hydrolysis. Subsequently, EF-G is released in complex with GDP and the ribosome translocates by 3 nts along the mRNA.^6–8^ The EF-G conformational change between pre-translocation and post-translocation states likely produces a power stroke that would be the driving force for translocation (Figure 1a). Thus, translocation mechanism concerns two key mechanical components: the power stroke produced by EF-G and the precise 3-nts displacement of the ribosome. An intriguing question is whether these two factors are correlated for maintaining the ribosome on the correct reading frame.^9^ To investigate this correlation, the Q508K and H584K mutants (*E. coli* EF-G sequence, numbered from the first methionine) will be utilized to determine how alterations in EF-G’s power stroke impact translocation efficiency and fidelity. These mutants were selected because structural and functional studies have shown they are highly conserved sites in domain IV of EF-G, potentially playing significant roles in maintaining mRNA reading frame. Some variations are shown to be lethal, drastically reducing the translocation kinetics and causing high-percentage frameshifting.^10, 11^

**Figure 1.**
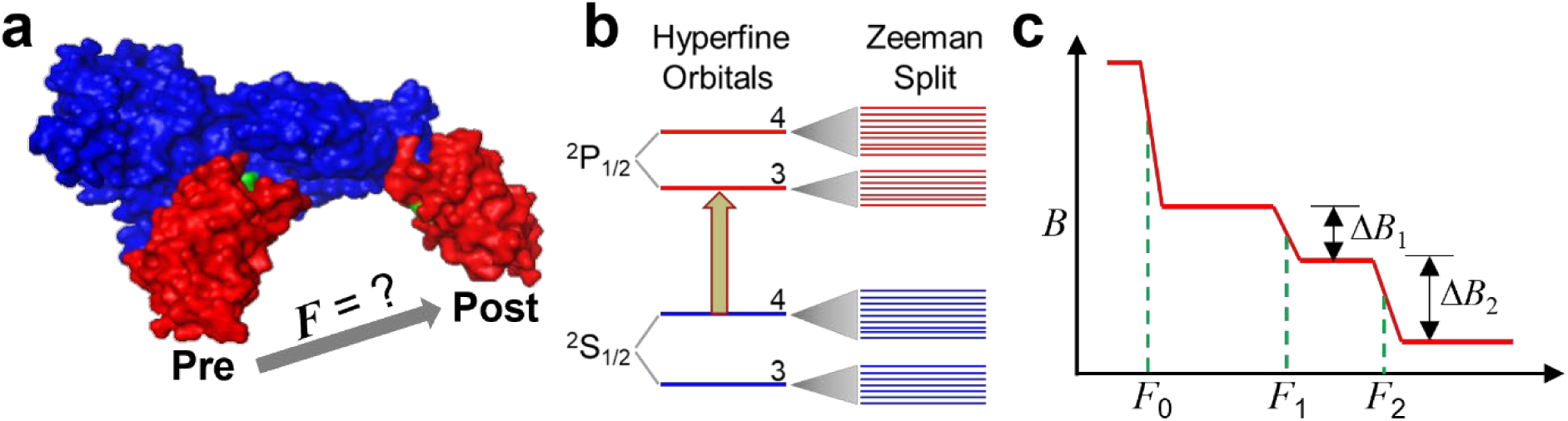
Quantum sensing techniques for EF-G power stroke and ribosome translocation. **a.** Conformational change of EF-G that probably produces power stroke *F*. Pre-translocation and post-translocation EF-G structures are overlaid. Red: domain IV; blue: the rest domains. **b.** Atomic magnetometry based on quantum coherence among the Zeeman levels in ^133^Cs produced by polarized excitation. First column: hyperfine orbitals after electron-nuclear spin coupling with numbers indicating the total quantum number; second column: Zeeman splitting of the orbitals in the presence of an external magnetic field. **c.** SURFS technique based on atomic magnetometry, in which biochemical bonds can be precisely resolved based on their dissociation forces. The changes in magnetic signals indicate the abundance of the bonds.

Quantifying EF-G power stroke has been a technical challenge. Three different experimental approaches have been reported in the literature.^12–14^ We first reported the power stroke of 89 pN using a series of DNA-mRNA duplexes as force gauges. We also showed that this force is directional, which is consistent with the ribosome’s moving direction and EF-G conformational changes.^12^ Liu *et al.* used optical tweezers to pull the ribosome-mRNA system, in which the ribosome was immobilized on the surface and mRNA was labeled with a bead controlled by the optical tweezers. A smaller power stroke of 13 pN was concluded.^13^ The results by Chen and co-workers using an indirect method favored the model with significant power stroke; however, no specific value was derived.^14^ We later demonstrated that the power stroke was reduced when EF-G was crosslinked to restrict its conformational change; power stroke could also be reduced by the binding of fusidic acid that constrains EF-G conformational changes.^15^ However, no correlation between power stroke and translocation fidelity has been observed, possibly because no alteration was attempted for the critical domain IV of EF-G. In addition, none of the previous force measurements have directly compared the power stroke to a well-defined mechanical force acting on the same ribosome complex. A more refined experimental approach is needed to enhance the quantification of force amplitude.

Recent developments in quantum sensing and force spectroscopy, among which are atomic magnetometry ^16–19^ and super-resolution force spectroscopy (SURFS), ^20^ have provided viable tools to both measure both power stroke and resolve ribosome translocation. Atomic magnetometers rely on the quantum coherence between Zeeman levels in vapor-phase alkali atoms to detect magnetic signals (Figure 1b). Sensitivity at the level of fT/(Hz)^1/2^ has been routinely achieved.^17^ In addition, a unique characteristic of atomic magnetometers is the long detection distance, which allows them to detect a much larger sample area compared to microscopic techniques. Based on atomic magnetometry, our groups invented SURFS by integrating acoustic force with magnetic sensing.^20, 21^ DNA probes labeled with magnetic beads form duplexes with the exposed segment of mRNA in the ribosome complex. Precise external forces were applied to dissociate the duplexes, which were measured as a decrease in the magnetic signal by an atomic magnetometer (Figure 1c). The nonspecific beads were first removed by a weak force *F*_0_ to ensure high-resolution force spectra. The ribosome translocation steps were revealed by the different dissociation forces, *F*_1_ and *F*_2_, of the DNA-mRNA duplexes in the pre-translocation and post-translocation complexes. The abundances of the complexes are indicated by the decrease of magnetic signals, Δ*B*_1_ and Δ*B*_2_, at the corresponding forces. We have achieved 0.4 pN force resolution,^21^ compared to the typical 10-20 pN force distribution by other force techniques that include atomic force spectroscopy, optical tweezers, magnetic tweezers, and acoustic force spectroscopy.^22–26^ The step size of ribosome translocation can be monitored with sub-nucleotide precision.^27, 28^ Our quantum-based techniques have successfully revealed “-1”, “-2”, and “+1” frameshifts that were consistent with conventional molecular biology techniques.^29, 30^

In this work, we report a unique combination of methods that measure power stroke with precise quantification and determine translocation steps from both ends of the ribosome, in the presence of mutated EF-Gs at two critical residues. Power stroke and acoustic force were applied sequentially onto the same ribosome systems, which yields precise percentages of DNA-mRNA force gauges dissociated by each force. The translocation steps were determined with single-nt resolution for both sides of the ribosome. We show direct evidence that a weaker power stroke by one mutated EF-G induced “-1” frameshift, whereas the other mutated EF-G generated the same power stroke as the wild type and did not induce frameshifting. The clear correlation between power stroke and translocation fidelity shows the important role of specific noncovalent bonds between EF-G domain IV and tRNA in maintaining the correct reading frame in protein synthesis.

## RESULTS AND DISCUSSION

### GTPase of EF-Gs

Two point mutations were performed on EF-G, Q508K and H584K, which were chosen due to their critical interactions with the A-site tRNA during translocation (Figure 2a).^31^ Structurally, the glutamine-to-lysine mutation, which is not studied in the literature, removes the carbonyl structure from the residue and increases the length by one carbon. Both residues are polar, which may allow the mutation to maintain its hydrogen bonding with A37 of the A-site tRNA. In contrast, the histidine-to-lysine mutation causes a significantly structural change, eliminating the heterocyclic imidazole ring, which may disrupt its specific interaction with residue 36 of the tRNA, the leading position of codon-anticodon helix during translocation.^11, 32^ The mutations were introduced using polymerase chain reaction (PCR) and purified using fast protein liquid chromatography (FPLC) (Supplementary Information, Scheme S1). Purified mutations were quantified using sodium dodecyl sulfate-polyacrylamide gel electrophoresis (SDS-PAGE) (Figure 2b, Supplementary Information, Figure S1), and sequencing confirmed the correct mutations (Supplementary Information, Figure S2).

**Figure 2.**
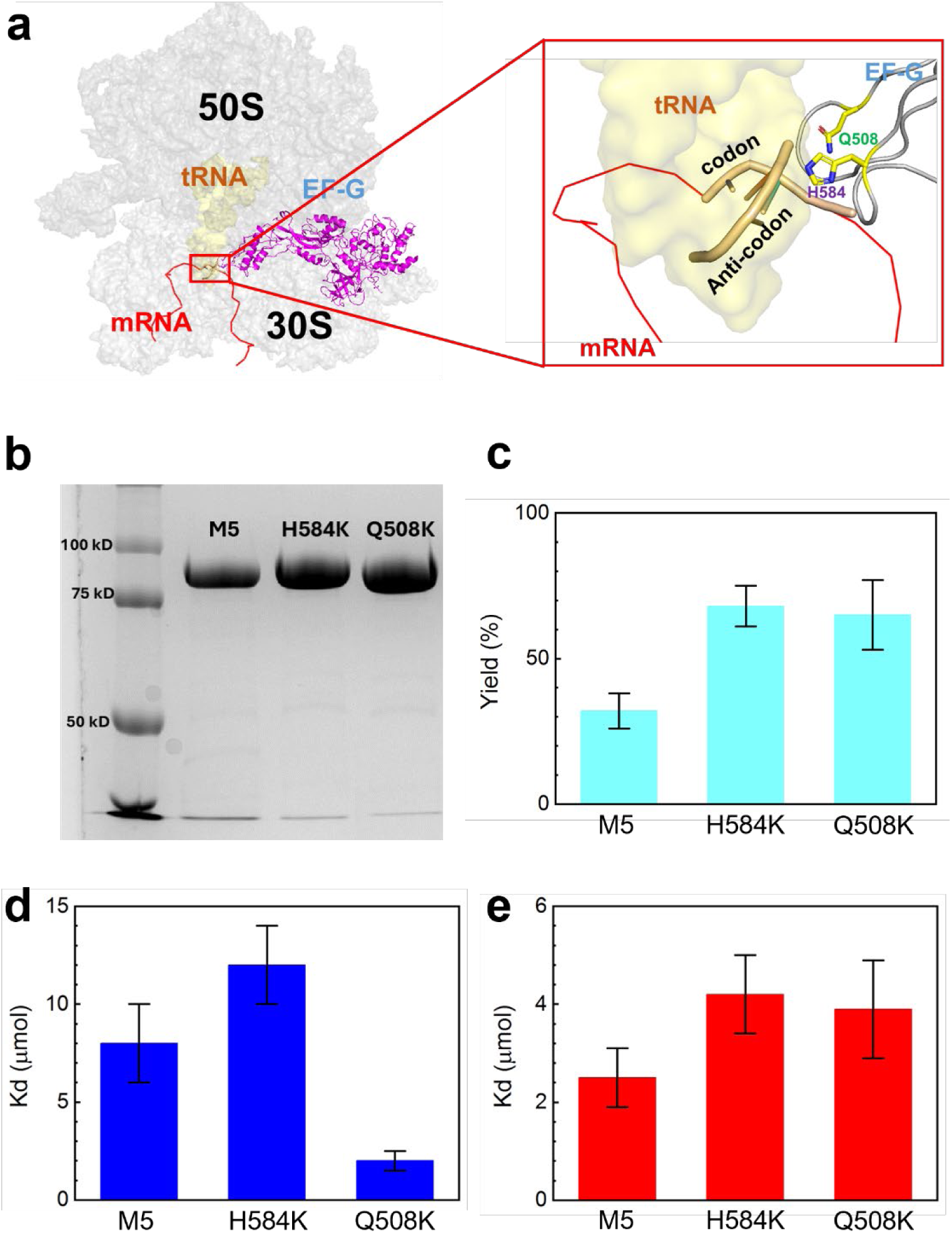
EF-G domain IV mutations and GTPase activity. **a.** Ribbon structure of ribosome, tRNA and EF-G (PDB: 4w29). The closeup view highlights H-bonding between tRNA and residues H584 or Q508, depicted in the red box. **b.** 8% SDS-PAGE gel of EF-G mutants following FPLC purification. **c.** GTP hydrolysis yields as determined by TLC. **d,e.** Mant-GTP binding and mant-GDP binding, respectively. K_d_ was calculated via a mant-GTP and mant-GDP titration procedure published previously.^33, 34^

GTPase assays were performed on mutated and nonmutated EF-G to determine the mutagenesis effects on protein activity. The unmutated EF-G, named as M5 in this work, served as the template for introducing mutations and has been shown to support normal translocation.^15^ Thin layer chromatography (TLC) assay was used to determine the overall percentages of fluorescently active mant-GTP hydrolyzed to mant-GDP (Figure 2c).^33^ The percentages were 68±7, 65±12, and 32±6 for H584K, Q508K, and M5, respectively, after 6 minutes. These results verified that both mutated EF-Gs exhibited hydrolysis activities on the same order of magnitude as M5 under our experimental conditions. Furthermore, dissociation constants for GTP and GDP binding (Supplementary Information, Figure S3) were determined using an established method from the literature.^34^ The values, plotted in Figures 2d and 2e, are within reasonable experimental variations compared to prior reports.^34–37^

### Power stroke of EF-Gs

Power stroke measurements are shown in Figure 3. A series of DNA probes, DNA_12 to DNA_19, were used to form 12-19 bp duplexes, respectively, with the 3′ end of the mRNA. Subsequently these duplexes were used as force rulers to gauge the EF-G power stroke. When the power stroke was greater than the dissociation force of the DNA-mRNA duplexes, a percentage of the duplexes would dissociate, indicated as a decrease in the magnetic signal.^15^ This is because of the randomization of the magnetic dipoles of the dissociated beads attached to the 5’ end of mRNA. To accurately determine the dissociation percentage, we applied acoustic force to dissociate all the remaining duplexes after EF-G’s power stroke (Figure 3a). Therefore, the dissociation due to power stroke can be calculated from dividing the magnetic signal decrease due to power stroke by the overall signal decrease after applying sonication. This new approach provides direct signal calibration compared to our previous work in which the percentages were normalized to the shortest force ruler, although the same power stroke results were obtained. The magnetic signal of the samples was measured by translating the sample to a downstream atomic magnetometer at ∼200 mm from the location of force application (Supplementary Information, Scheme S2). The sensitivity was approximately 300 fT/Hz^1/2^.^38^

**Figure 3.**
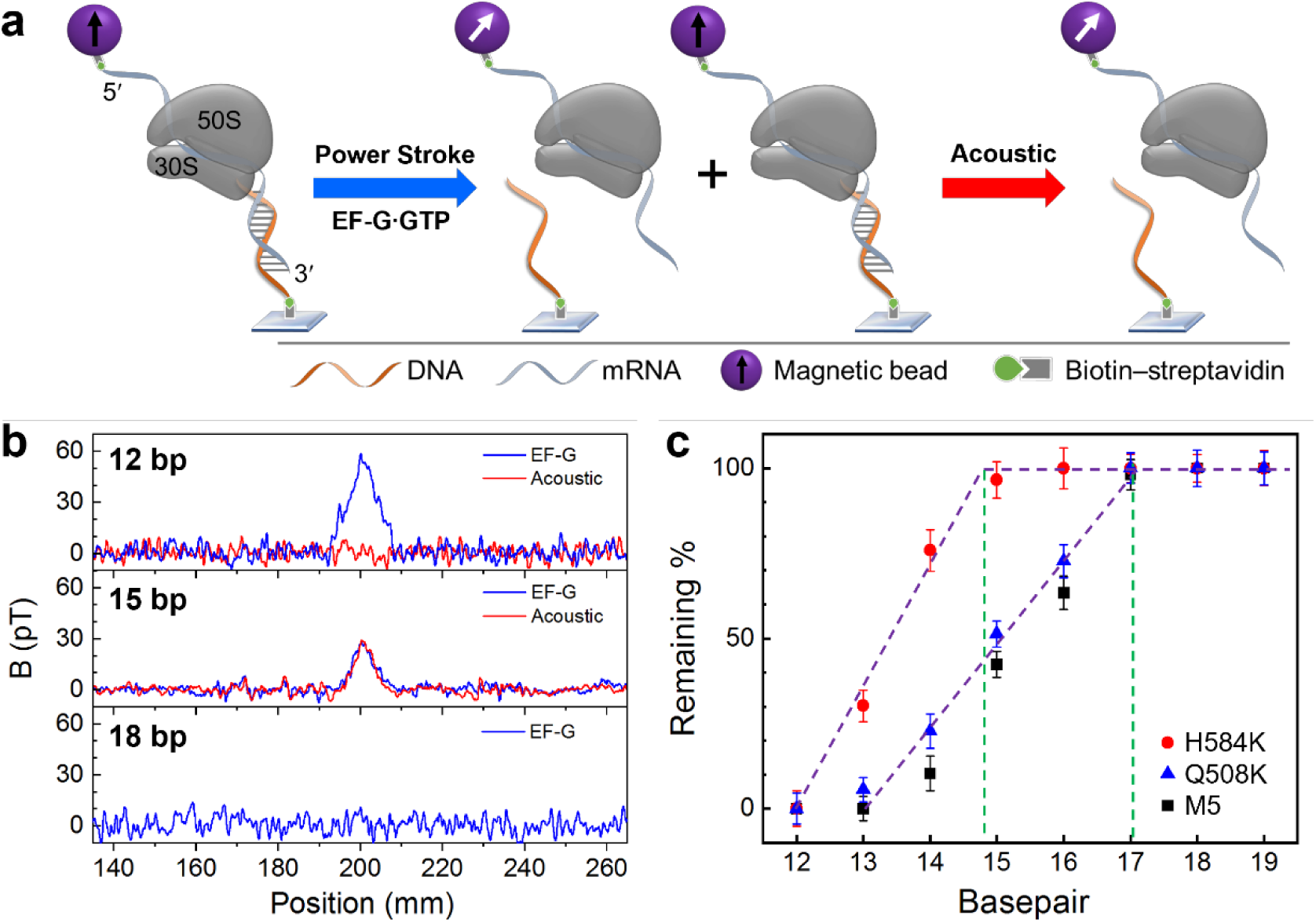
Power stroke of EF-Gs. **a.** Measuring method. The ribosome complex forms mRNA-DNA duplex at the 3′ end and is labeled with a magnetic bead at the 5′ end. EF-G power stroke will first induce duplex dissociation, and then the remaining duplex will then be dissociated by acoustic force. **b.** Magnetic signal changes for three different bps under EF-G power stroke and acoustic force. **c.** Plots of remaining duplexes for duplexes of 12-19 bps, in the presence of EF-Gs. The green lines indicate the onset of the dissociated duplex by EF-Gs.

Figure 3b shows three typical magnetic signal changes for rulers at three different lengths and under two different forces. In the top panel for 12 bp duplexes, a large decrease of 59 pT was observed by EF-G power stroke and no additional decrease was observed for acoustic force. For 15 bp duplexes (middle panel), the signal decrease due to EF-G is 30 pT, indicating that EF-G power stroke exceeds the dissociation force of the duplex; further applying acoustic radiation force produced another signal decrease, at approximately 30 pT. This means the overall yield of the duplex was 60 pT, out of which 50% were dissociated by EF-G power stroke. The last case is for 18 bp duplexes, in which no dissociation was observed by EF-G (bottom panel). Therefore, the dissociation percentage is zero and there is no need for applying acoustic force.

Using this method, we plotted the dissociation percentages for 12-19 bp duplexes in the presence of three different EF-Gs (Figure 3c). For both M5 and Q508K, the onset of dissociation occurred at 17 bp, which are consistent with our previous works.^12, 15^ Therefore, the power stroke for both EF-Gs was 89±11 pN. For H584K, however, the onset was at approximately 15 bp (14.8 bp for the intercept for the red plot on Figure 3c). Given the dissociation force for 14 and 15 bp being 50.1 and 62.0 pN, respectively^38^ (Supplementary Information, Figure S4), H584K’s power stroke was determined to be 60±6 pN. This amplitude is similar to that of EF-G crosslinked by a short molecular linker or EF-G bound with fusidic acid.^15^

### Ribosome translocation and frameshifting

To reveal potential differences in translocation fidelity for the mutated EF-Gs, we used the SURFS technique to probe the translocation steps from both 5′ and 3′ ends of the mRNA with single-nt resolution, which has been well established in our groups.^27, 39^ Figure 4a illustrates the detection scheme. The ribosome pre-translocation complex (Pre) was immobilized on the surface via the biotin on the 5′ end of the mRNA. A DNA probe was then incubated with the Pre complex to form duplexes with the exposed mRNA, at either the 5′ or 3′ end. For the 5′ end, the DNA probe forms 12 bp duplexes with dissociation force of 21 pN, and would form 15-bp duplex with the Post complex if the ribosome undergoes normal translocation with dissociation force of 52 pN. If the ribosome translocates only two nucleotides, 14 bp duplexes will be formed, which will be indicated by a weaker force of 45 pN. With force resolution of ∼2 pN in this work, we can easily distinguish the translocation steps of the ribosome. Similarly for the 3′ end, 14 bp duplexes will be formed between the DNA probe and Pre, and 11 bp for Post for normal translocation but 12 bp for “-1” frameshifting (Supplementary Information, Figure S3). Results in Figure 4b showed that the ribosome underwent normal translocation in the presence of Q508K or M5, in which it translocated by 3 nts from both the 5′ and 3′ ends of the mRNA. This correlates well with the normal power stroke of Q508K compared to M5. However, for H584K, which was only able to produce a much reduced power stroke, the ribosome translocated by 2 nts, i.e., “-1” frameshifting occurred. This result is consistent with previous studies on H584 mutation.^10, 11^ It is also interesting that the ribosome has the same displacements at the 5′ and 3′ ends. In other words, the mRNA threaded through the ribosome smoothly; no looped intermediate state was formed. These observations strongly suggest that EF-G power stroke is important for the ribosome to maintain the reading frame during translocation.

**Figure 4.**
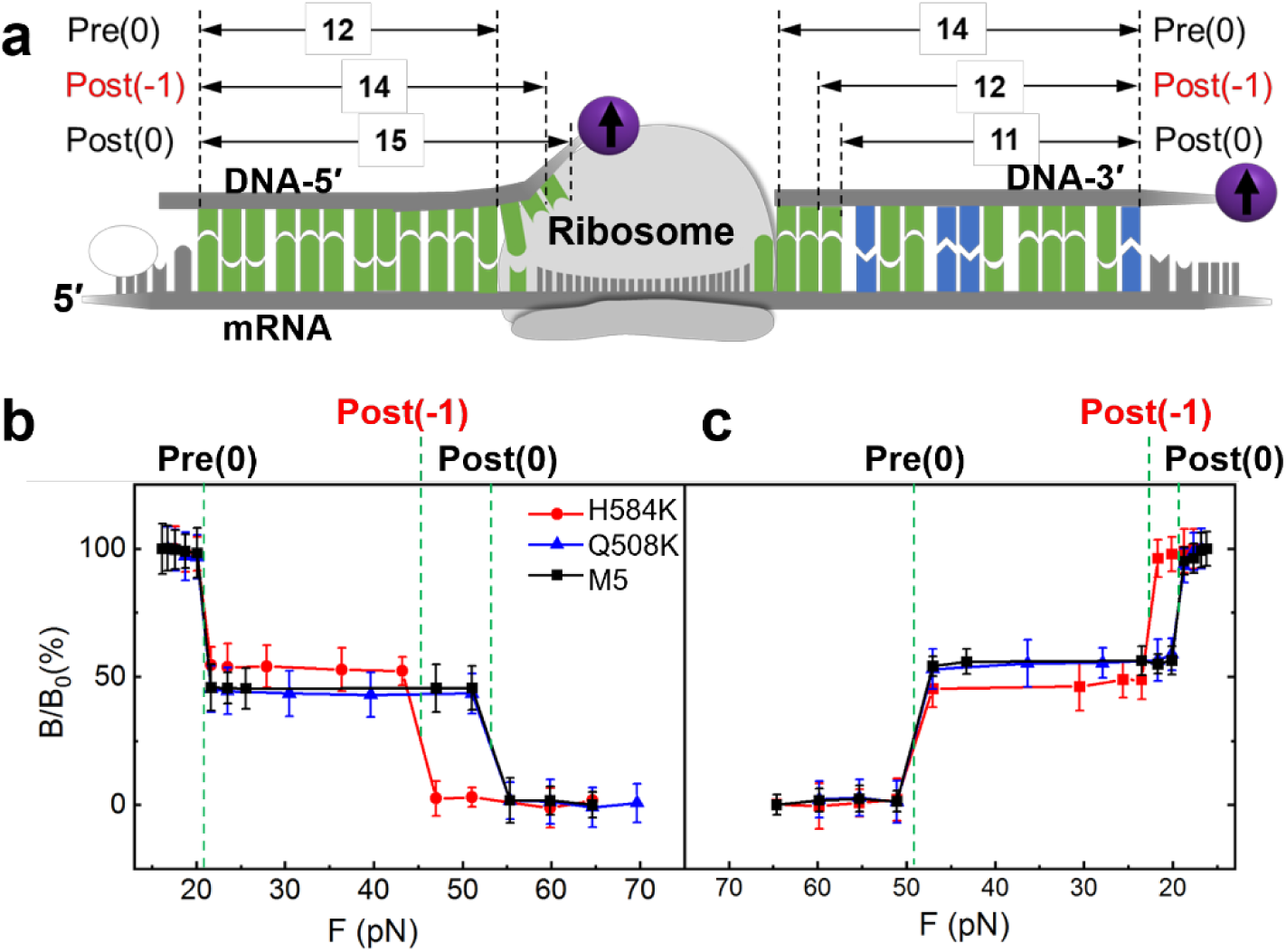
Translocation by EF-Gs. **a.** Probing scheme. The ribosome pre-translocation complex (Pre) was immobilized on the surface. DNA-5′ and DNA-3′ were used to probe ribosome motion on the 5′- and 3′-end of the mRNA, respectively. Pre(0) and Post(0) are the normal reading frame; Post(-1) is for “-1” frameshift in which the ribosome moved by 2 nts. The numbers with arrows indicate duplex bps between DNA probes and the exposed mRNA segments in the ribosome complex. **b,c.** Force spectra for the 5′-end and 3′-end, respectively. The transitions were assigned to different ribosome states based on the force values.

The clear correlation between a weaker power stroke and a reduced translocation step provides the first mechanical evidence demonstrating the vital role of specific interactions between EF-G and tRNA. A strong coupling allows EF-G to drive the full 3-nt translocation, while alteration of these interactions causes EF-G to be less effective. Consequently, translation errors such as “-1” frameshifting may occur (Figure 5). Structural studies have shown that EF-G interacts with A-site tRNA via H584 and Q508 residues by noncovalently bonding with tRNA’s U36 and A37 nucleotides, respectively.^10, 11, 32^ The H584K mutation likely altered the orientation of its interaction with the leading base pair of the minihelix along the translocation direction (Figure 5). Consequently, changes in force projection on the mRNA moving trajectory and the pivotal point on tRNA would reduce the power stroke’s effect on the activation energy barrier, according to the Bell’s model.^12^ Applying the same Bell’s model, the deceleration factor is approximately 40-folds for the 30 pN force decrease we observed at 25 °C, which is close to the 100-fold deceleration effect reported earlier.^40^ If changes in the pivotal point further increase the transition state distance, this effect could be even greater. In contrast, Q508K mutation was less likely to generate significant structural changes. Therefore, we propose that the power stroke likely plays multifaceted roles in regulating translocation, influencing not only through its amplitude but also via atomic-level interactions with the ribosome complex. The previously observed decrease in kinetics for H584K was attributed to increased frameshifting yield.^10, 11^ Our results here imply that this effect is likely due to altered force projection and a shifted pivotal point, leading to a less-reduced activation barrier and consequently slower translocation.

**Figure 5.**
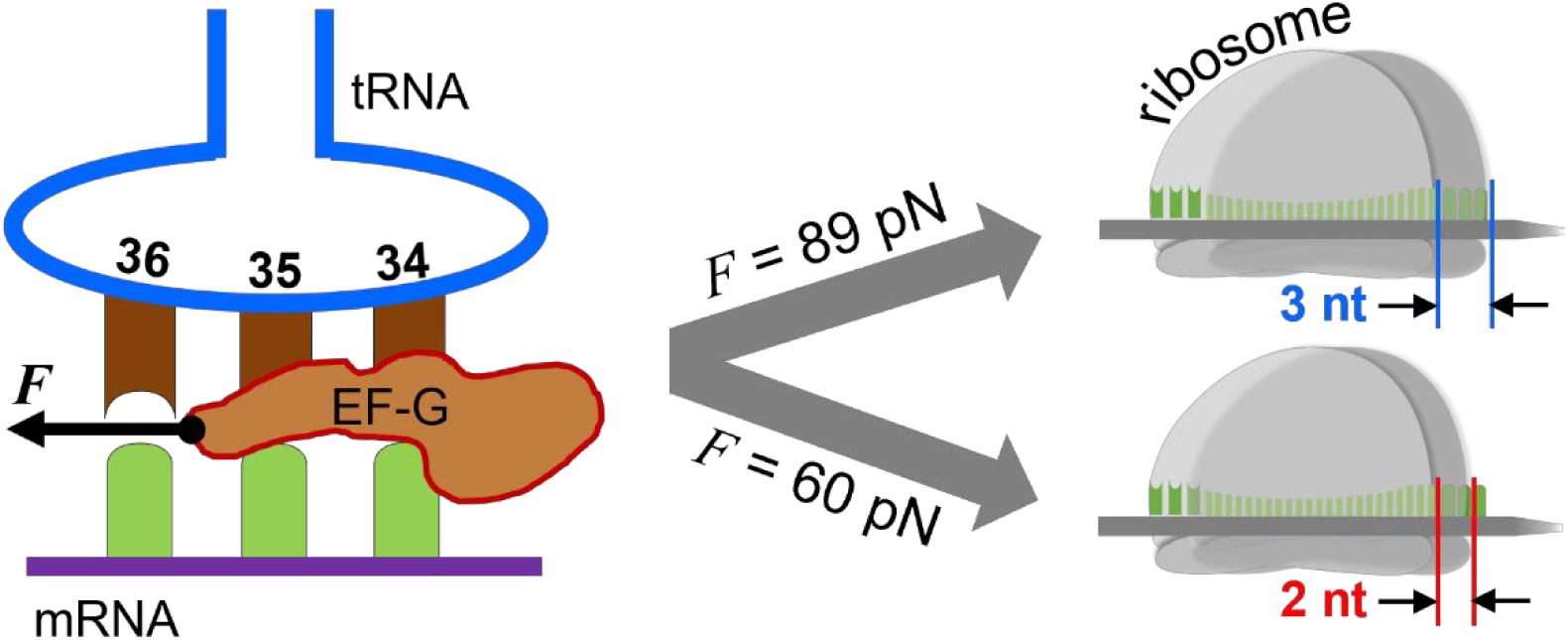
Altering EF-G–tRNA bonding may cause translation error. *F* indicates the power stroke generated by the EF-Gs (left). When the force is high, such as in the cases of M5 and Q508K, it leads to the correct reading frame after translocation, in which the ribosome translocates 3 nts on mRNA from both sides (right, top case). In contrast, a weakened power stroke by H584K correlates with “-1” frameshift in which the ribosome moved only 2 nts on both sides of the mRNA (right, bottom case).

## CONCLUSIONS

Our quantum-based techniques to probe power stroke and ribosome displacement offer a unique perspective for investigating complex biomolecular functions. The combination of magnetic labeling, DNA force gauges, and external mechanical force provides a viable approach to precisely determine the mechanical forces involved in protein conformational changes, which are difficult to be directly measured by optical- or fluorescence-based techniques. The application of atomic magnetometry, SURFS, and DNA displacement rulers provides sub-nt resolution for molecular motion, which gives precise local information during biomolecular functions that complement techniques for global structures and force measurements. This work shows that the integration of quantum sensing and force spectroscopy can be unique tools to investigate the mechanical aspects of biomolecular machines as complicated as the ribosome, where other techniques may have relatively limited viability.

## EXPERIMENTAL METHODS

### EF-G mutation and characterization

Site-directed mutagenesis of *Escherichia coli* was performed using a Q5^®^ High-Fidelity DNA Polymerase kit (New England Biolabs, NEB) via polymerase chain reaction (PCR). Forward and reverse primers were designed using NEB Base Changer and purchased from Integrated DNA Technologies (IDT). The completed PCR reaction product was transformed into MAX Efficiency^®^ DH5α™-T1R competent cells (ThermoFisher Scientific) and grown on Lysogeny Broth-Kanamycin (LB-Kan) agar plates overnight at 37 ℃. pDNA from colonies was isolated using an PureLink™ HQ Mini Plasmid DNA Purification Kit (ThermoFisher Scientific) and sequenced via Sanger sequencing. Sequenced mutants were transformed into One Shot™ BL21(DE3) cells (ThermoFisher Scientific) and grown in 3 liters of LB-Kan liquid medium until cultures entered log phase. The cultures were then induced using 1 M isopropyl ß-D-1-thiogalactopyranoside (IPTG) and incubated for three hours at 37 °C while shaking. Cells were lysed using lysozyme, DNase I, and bacterial extraction reagent (0.7% (w/v) n-octyl-1-thio-beta-D-glucopyranoside, 0.2% (w/v) Nonidet P-40, 0.1% (w/v) Triton^TM^ X-100, 50 mM Tris-Acetate (pH 7.60), 1 mM MgCl_2_, 0.5 mM CaCl_2_, and 0.1 mM EDTA Sodium Salt (pH 8.00)). Cell debris was pelleted, and the cell lysate was cycled through a 5 mL HisTrap^TM^ HP His tag protein purification column (Cytiva Life Sciences) using a Pharmacia LKB P-1 pump for three hours. The cycled Ni^2+^ column was connected to a ÄKTApurifier Fast Protein Liquid Chromatograph (FPLC) system, and the column was flushed with protein lysis buffer (50 mM NaH_2_PO_4_, 150 mM NaCl, and 4 mM BME, all from Sigma-Aldrich, pH 8.0). Protein was eluted using a gradient of protein elution buffer (50 mM NaH_2_PO_4_, 150 mM NaCl, 4 mM BME, 1 M imidazole (Sigma-Aldrich), pH 8.0). FPLC fractions were quantified via SDS-PAGE and fractions containing purified EF-G were stored in protein storage buffer (20 mM Tris-HCl, 10 mM MgCl_2_, 40 mM KCl, 4 mM BME, pH=7.5) at - 80 °C.

### Preparation of the ribosome complex

The 70S ribosomes were purified from *E. coli* MRE600 as previously described. The culture of *E. coli* MRE600 was propagated in LB medium in a shaker incubator at 37 °C and 200 rpm (revolution per minute) until optical density at 600 nm reached value of 0.6. The cells were harvested by centrifugation at 3,000×*g*, 4 °C, for 20 minutes. The cell pellet was washed with cold buffer I (50 mM Tris-HCl, pH 7.6, 10 mM MgCl_2_, 100 mM NH_4_Cl, 6 mM BME, 0.5 mM EDTA). Then the cells were resuspended in the same buffer with 0.2 mg/mL egg white lysozyme and 2 μg/mL DNase I and incubated for 30 min on ice. Afterward the cells were lysed by sonication for 5 minutes (10 s pulse, 20 s pause) on ice. Cell debris were removed by two consecutive centrifugations at 14,500×*g*, 4 °C, for 1 hour each. Cleared supernatant was layered on top of pre-cooled 1.1 M sucrose cushion in buffer II (20 mM Tris-HCl, pH 7.6, 10 mM MgCl_2_, 500 mM NH_4_Cl, 6 mM BME, 0.5 mM EDTA) with volume ratio 1:1, respectively. The ribosomes were pelleted by centrifugation on Beckman XL-80 ultracentrifuge with Type 45 Ti Fixed-Angle rotor at 120,000×*g*, 4 °C, for 20 hours. The collected pellet was carefully rinsed with buffer I, then the ribosomes were resuspended in the same buffer, and the concentration of NH_4_Cl was adjusted to 400 mM. The ribosomes were pelleted again by ultracentrifugation at 120,000×*g*, 4 °C, for 20 hours. The pellet was rinsed with buffer I and finally resuspended in minimum volume of the same buffer. The ribosome concentration was determined by UV absorbance measurement at 260 nm using conversion coefficient 1 A260 unit = 23 pmol (10^-12^ mol) of 70S ribosomes. The prepared ribosome solution was aliquoted, flash-frozen in liquid nitrogen, and stored at -80 °C.

All the mixtures were in TAM_10_ buffer, which consisted of 20 mM Tris-HCl (pH 7.5), 10 mM MgCl_2_, 30 mM NH_4_Cl, 70 mM KCl, 0.5 mM EDTA, and 7 mM BME (2-mercaptoethanol). Three mixtures were prepared: the ribosome mix, Tu0G mix, Phe mix. The ribosome mix contains 1 μM ribosome, 1.5 μM each of initiation factors IF1, IF2, IF3, 2 μM of mRNA, 4 μM of charged fMet-tRNA^fMet^, and 4 mM of GTP. The Tu0G mix contained 4 μM EF-Tu, 4 mM GTP, 4 mM 2-phosphoenolpyruvate (PEP), and 0.02 mg/mL Pyruvate Kinase. The Phe mix contained 100 mM Tris (pH 7.8), 20 mM MgCl_2_, 1 mM EDTA, 4 mM ATP, 7 mM BME, 2 μM tRNA^Phe^ synthetase (PheRS), 2 A_260_ units/mL tRNA^Phe^, and 0.25 mM phenylalanine. The ribosome mix, Tu0G mix, and Phe mix were combined in a volume ratio of 1:2:2, then incubated at room temperature for 2 minutes. The resulting MF-Pre ribosome complex was added on top of 1.1 M sucrose cushion and purified by centrifugation with a Hitachi CS150FNX ultracentrifuge (S140AT rotor, 200,000-400,000×*g*, 4 ℃, 3 hours).

Additional details regarding sample preparation are available in one of our recent publications.^41^ The viability of our ribosome complex for translocation and frameshifting has been demonstrated in our many previous publications using optical, magnetic, and other molecular biology assays.^29, 30, 41^

### Power stroke measurements

The sample wells were of 4×3 mm^2^ area formed by glueing a piece of biotin-coated glass with a plastic holder (2 mm thick). After incubating with 0.25 mg/mL streptavidin for 50 minutes, the extra streptavidin was rinsed twice with water. The probing DNA strand was immobilized on the surface and incubated for 1 hour. After rinsing once with TAM10 buffer, the MF-Pre ribosome complex was added and incubated overnight to form duplexes with DNA. Afterward, the magnetic beads were added into the sample well and incubated for 2 hours. The sample was magnetized for 2 minutes using a permanent magnet (∼0.5 T) and centrifuged (5427R, Eppendorf) at 820×*g* for 20 minutes to remove non-specifically bound magnetic particles. The initial magnetic signal of the samples was measured by a home-built atomic magnetometer. The EF-G was added and incubated with duplexes for 1hr. The magnetic signal of the samples was measured again before and after applied acoustic force. The total magnetic signal was obtained by the magnetic signal before EF-G subtracting the signal after the acoustic force. The remnant magnetic signal was obtained by the magnetic signal after EF-G subtracting the signal after the acoustic force. Percentages of remnant magnetic beads were obtained by dividing the remnant magnetic signal by the total magnetic signal.

The mRNA sequence is 5′-Bio/ C AAC UGU UAA UUA AAU UAA AUU AAA AAG GAA AUA AAA AUG UUU GAA AA<u>A</u> <u>AAG</u> <u>UAC</u> <u>GUA</u> <u>AAU</u> <u>CUA</u> <u>CUG</u> CUG AAC UC-3′. The underscored nucleotides will form 12-19 bp duplexes with the DNA force rulers depending on the lengths of the respective rulers. It was purchased from Integrated DNA Technologies (IDT) with HPLC purification.

The DNA rulers are:

PSNew_12: 5′-/BioTEG/CT CAA GTC GTC ATC TTT TAC GTA CTT T-3′

PSNew_13: 5′-/BioTEG/CT CAA GTC GTC ATC ATT TAC GTA CTT T-3′

PSNew_14: 5′-/BioTEG/CT CAA GTC GTC ATG ATT TAC GTA CTT T-3′

PSNew_15: 5′-/BioTEG/CT CAA GTC GTC AAG ATT TAC GTA CTT T-3′

PSNew_16: 5′-/BioTEG/CT CAA GTC GTC TAG ATT TAC GTA CTT T-3′

PSNew_17: 5′-/BioTEG/CT CAA GTC GTG TAG ATT TAC GTA CTT T-3′

PSNew_18: 5′-/BioTEG/CT CAA GTC GAG TAG ATT TAC GTA CTT T-3′

PSNew_19: 5′-/BioTEG/CT CAA GTC CAG TAG ATT TAC GTA CTT T-3′

The underscored nucleotides are complementary to the mRNA sequence. All DNA oligomers were purchased from IDT with HPLC purification.

### Ribosome translocation and frameshifting

The sample wells were the same as in the power stroke experiments. The difference is that the MF-Pre ribosome complex was immobilized on the surface via the 5′-end biotin on the mRNA by incubating for 1 hour. In the case of post complex, 1 µM Pre complex was incubated with 2 µM EF-G, 4 mM GTP, 4 mM PEP, and 0.02 mg/mL pyruvate kinase at 37 °C for 30 minutes. Then, 1 μM probing DNA was added and incubated overnight to form duplexes with the uncovered mRNA in the ribosome complex. Afterward, streptavidin-coated magnetic beads (M280) were introduced into the sample well and incubated for 2 hours. Before the force measurements, the sample was magnetized for 2 minutes using a permanent magnet (∼0.5 T) and centrifuged (5427R, Eppendorf) at 820×*g* for 20 minutes to remove non-specifically bound magnetic particles.

The sequence for the 3′ DNA probe is 5′-/BioTEG/CTC AAG TCG TCA T*GA TTT ACG TAC TTT*-3′, which forms 14 bp duplex with pre-translocation ribosome complex. The sequence for the 5′ DNA probe is 5′-/BioTEG/T_50_ AAA TTA *ATT TAA TTT AAT*-3′, which forms 12 bp duplex with the pre-translocation ribosome complex.

## Supporting information

Supplemental schemes and figures

## Acknowledgements

This work was supported by the US National Institutes of Health (R01GM111452, Y.W., S.X.) and National Science Foundation (2130427, S.X., Y.W.). J.H.S. was supported by US National Institutes of Health (3R01GM111452-07S1).

## Author contributions

Y.W. and S.X. designed the overall project. J.H.S. and Y.W. prepared the samples and performed the biological assays. Y.C. performed force measurements and translocation. All authors contributed to data analysis, discussion, and preparation of the manuscript.

## Conflicts of interest

S.X. and Y. W. declare financial interest in UForce Biotechnology, LLC.

## Data Availability

Additional information and data are available in the online ESI. The datasets generated and/or analyzed during the current study are available in the UniProt repository, SPIN200033120 and SPIN200033125.

## Notes

### Summary of Updates

We have revised the figures and corrected the conflict of interest statement.

