## Supplemental schemes and figures for "EF-G Mutations Reveal Correlation between Power Stroke and Translocation Fidelity in Protein Synthesis"

**Table of Contents**

1. Flowchart of the EF-G mutation, with *Scheme S1*;
2. SDS-PAGE gel photo of the mutated EF-Gs, with *Figure S1*;
3. Sequencing results of the mutated EF-Gs, with *Figure S2*;
4. GTP and GDP binding assay, with *Figure S3*;
5. Magnetic detection with optically pumped magnetometer, with *Scheme S2*;
6. Force calibration for the DNA-mRNA duplexes, with *Figure S4*.

1. Flowchart of EF-G mutation

The preparation procedure is summarized in the following scheme.

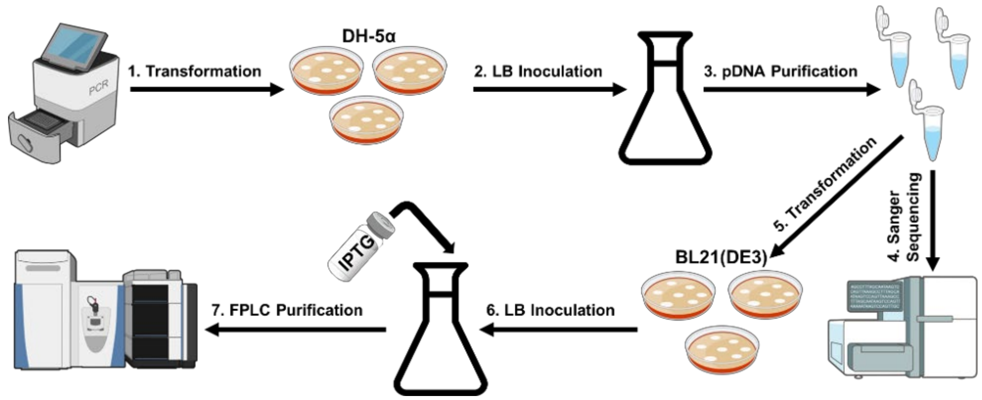


**Scheme S1.** EF-G mutagenesis flowchart.

2. SDS-PAGE gel photo of the mutated EF-Gs

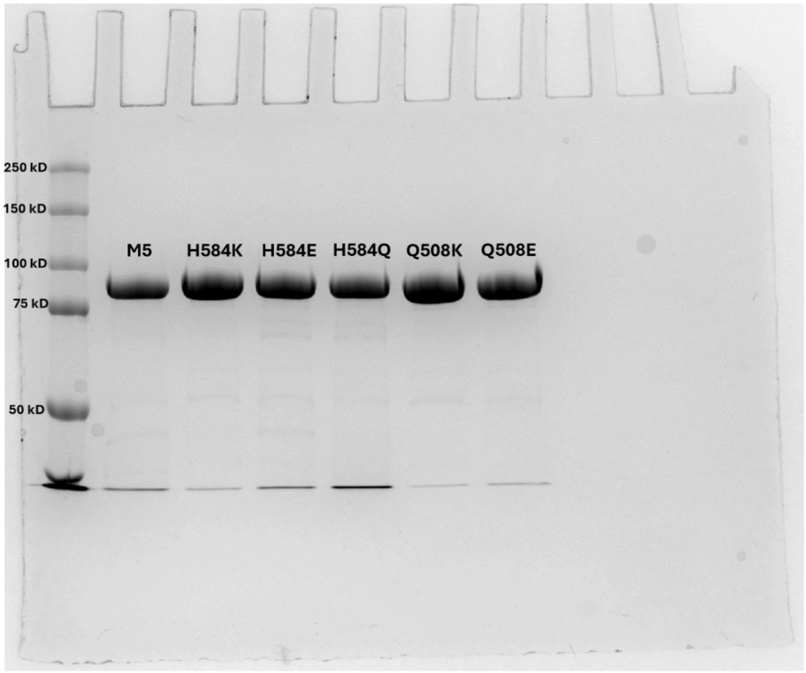


**Figure S1.** SDS-PAGE gel photo of five mutated EF-Gs. Two of the mutations, H584K and Q508K, are reported in this work.

3. Sequencing results of the mutated EF-Gs

The mutated EF-Gs were confirmed using Sanger sequencing (Figure S2) (Epoch Life Sciences). Sequencing was performed using T7 (5′–TAA TAC GAC TCA CTA TAG GG –3′), P2 (5′–GGT TCC GCT GCA GCT GG –3′), P3 (5′–CTC CGT GAA AGC TGC AC –3′), and P4 (5′–ACG AGT TCA TCA ACG AC –3′) primers. No offsite mutations or frameshifting was present due to non-selective forward and reverse primer binding during PCR.

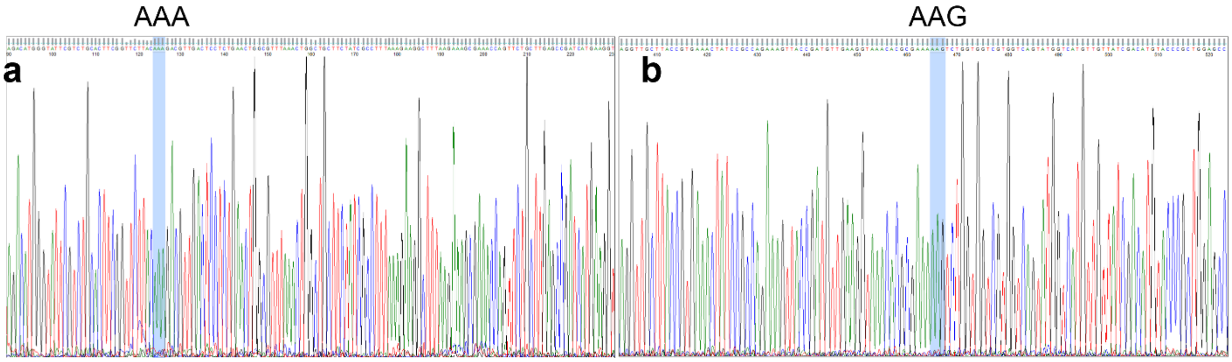


**Figure S2.** Sequencing results for **a**) H584K and **b**) Q508K. The new residue K is indicated by its genetic code AAA or AAG.

4. GTP and GDP binding assay

Mant-GXP (mant: 2′/3′-*O*-*N*-Methylanthraniloyl, X = T or D, both from Sigma Aldrich) binding assays were performed on EF-G M5 and domain IV mutant EF-G to determine the effects mutations have on GXP binding and dissociation. Both mant-GXP and EF-G solutions were prepared in TAM10 buffer with BME. The EF-G samples were then diluted with 40 µL of 36 µM Mant-GXP. The final concentrations of the samples were 32, 16, 8, 4, 2, and 0 µM following the addition of the mant-GXP solution. The final mant-GXP concentration was 16 µM following dilution. A blank sample was also made using only 50 µL of TAM10 buffer with BME. 40 µL of each concentration and blank was transferred to fluorescence tubes and the fluorescence measurements were performed three times for each sample. The averaged values were fitted using the following equation to obtain binding affinity $K$:

$$y = 16A + \frac{B - A - D}{2}\left[(x + 16 + K) - \sqrt{|(x + 16 + K)^2 - 64x|}\right] + Dx + C$$

Here, $A$ is mant fluorescence without EF-G, $B$ is fluorescence of EF-G/mant-GXP to be fitted, $C$ is offset for background, and $D$ is fluorescence of EF-G only. For GTP, A = 2397.15, C= 133.76, D = 8.88; for GDP: A =1966.68, C = 130.84, D = 8.88.

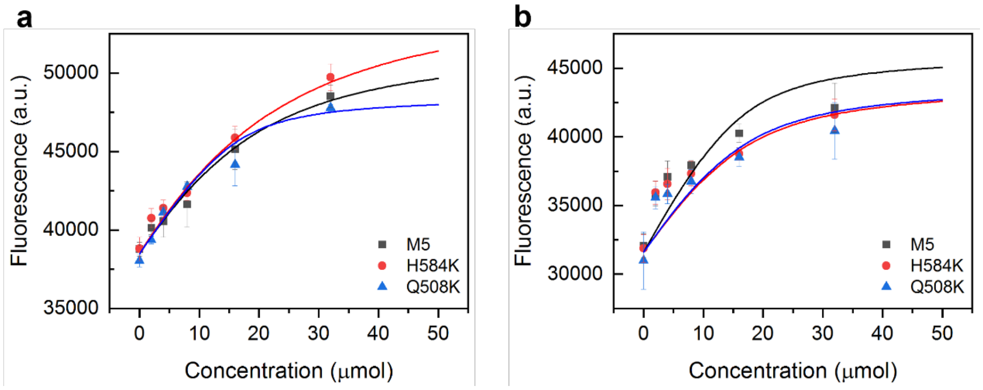


**Figure S3.** Determination of EF-G binding affinity with **a**) mant-GTP and **b**) mant-GDP.

5. Magnetic detection with optically pumped magnetometer

The following scheme shows the main components of magnetic detection using an atomic magnetometer.

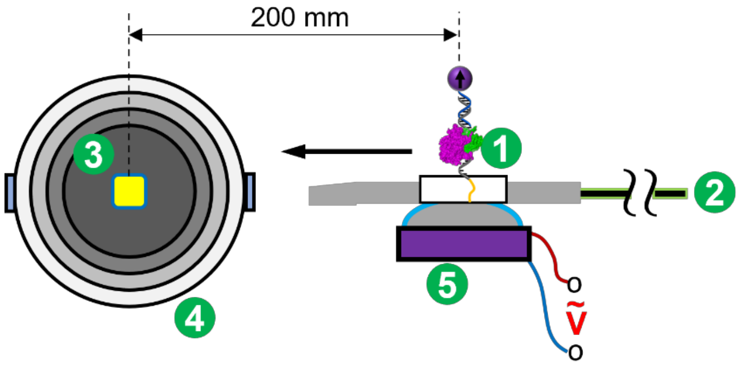

**Scheme S2.** Scanning magnetic detection with an optically pumped magnetometer. 1, sample with magnetically labeled ribosome system; 2, linear motor; 3, atomic sensor; 4, four-layer magnetic shield; 5, piezo to produce acoustic force.

6. Force calibration for the DNA-mRNA duplexes

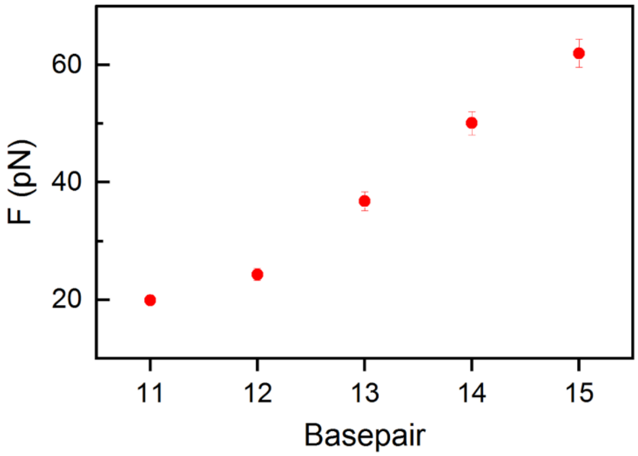


**Figure S4.** Force calibration for the DNA-mRNA duplexes.
